# Transcription factor HSFA7b controls ethylene signaling and meristem maintenance at the shoot apical meristem during thermomemory

**DOI:** 10.1101/2022.10.26.513826

**Authors:** Sheeba John, Federico Apelt, Amit Kumar, Dominik Bents, Maria Grazia Annunziata, Franziska Fichtner, Bernd Mueller-Roeber, Justyna J. Olas

## Abstract

The shoot apical meristem (SAM) is responsible for overall shoot growth by generating all above-ground structures. Recent research identified that the SAM displays an autonomous heat stress (HS) memory of a previous non-lethal HS event. Considering the importance of the SAM for plant growth it is essential to unlock how its thermomemory is mechanistically controlled. Here, we report that *HEAT SHOCK TRANSCRIPTION FACTOR A7b* (*HSFA7b*) plays a crucial role in this process in Arabidopsis. We found that HSFA7b directly regulates ethylene response at the SAM by binding to promoters of key regulators of ethylene signaling including *ETHYLENE-INSENSITIVE 3* to establish thermotolerance. Moreover, HSFA7b controls maintenance of the SAM stem cell pool during thermomemory by regulating the expression of the master regulator *WUSCHEL* through direct transcriptional activation of the *SPLAYED* chromatin remodelling factor.

## Introduction

As sessile organisms plants are frequently exposed to unpredictable and often life-threatening environmental challenges like e.g., heat stress (HS); however, evolution has established response strategies to cope with those variable conditions to ensure survival. In particular, plants employ a stress memory mechanism allowing them to ‘memorize’ the exposure to a first, moderate and non-lethal stress (called priming), during which information about the past stress is stored (memory phase) to facilitate a faster or stronger response to the next potentially more threatening stress signal (triggering)^1^. The initial phase of a HS response (priming and memory) involves a complex interplay between transcription factors (TFs) of the HEAT SHOCK FACTOR (HSF) family and HS memory genes^2,3^. HSFs bind to heat shock elements (HSEs, with a conserved 5’-nGAAnnTTCn-3 ‘ sequence) in promoters of HS-inducible genes including *HEAT SHOCK PROTEIN (HSP*) encoding chaperones^4,5^ to confer thermotolerance. Although components of the HS memory machinery have been characterized^6^, our understanding of the mechanisms underlying HS stress memory is scarce. To date, molecular HS memory has mainly been studied in whole *Arabidopsis thaliana* (Arabidopsis) plants^7–9^. However, it was recently shown that the shoot apical meristem (SAM) of Arabidopsis can directly sense changes in temperature and display a strong transcriptional HS memory for genes involved in protein folding, primary carbohydrate metabolism, and meristem maintenance^10^. Importantly, the SAM engages a HS response and memory network that in many aspects differs from that of whole seedlings: the expressional timing of HS memory genes and the types of genes generating thermomemory, including HSFs, are different between organs, providing strong evidence for tissue- or organ-specific heat memories.

The SAM represents a highly organized assortment of cells required for proper and continuous above-ground growth of plants^11^. It harbors stem cells whose descendants form shoot structures like leaves, flowers and derivatives thereof (seeds and fruits)^12^. SAM maintenance is controlled by a negative feedback loop between CLAVATA 3 (CLV3) and WUSCHEL (WUS)^13^ whereby WUS protein controls stem cell fate by activating *CLV3* transcription, while CLV3-mediated signaling limits stem cell accumulation by restricting *WUS* expression^13^. The balance between stem cell loss and renewal allows plants to form new organs throughout their entire lifespan^12^, supporting the plant’s high developmental flexibility. Apart from CLV3, other factors, including chromatin remodeling actors, affect *WUS* expression^14^. For example, SPLAYED (SYD), an SNF2 chromatin-remodeling ATPase, controls *WUS* transcription by binding to *cis*-elements in its promoter^15^.

Surprisingly little is currently known about how SAM maintenance genes are transcriptionally controlled by stress. Considering the key importance of the SAM for plant growth we expect dedicated molecular mechanisms allowing the SAM to respond appropriately to environmental challenges. This is particularly relevant for seedlings that have not yet established axillary meristems; a severe damage of the SAM by abiotic stress will lead to termination of shoot growth and, hence, death of the individual, precluding inheritance of genetic information to the next generation.

Here, we report that *HSFA7a* and *HSFA7b* are involved in HS memory at the SAM. We show that HS memory-related genes at the SAM are under direct transcriptional control of HSFA7b, and demonstrate that HSFA7b regulates ethylene response during HS by binding to HSEs in the promoter regions of ethylene-related genes. Furthermore, we find that *WUS* displays ectopic expression at the SAM of wild-type Arabidopsis plants in response to thermopriming. Our analyses revealed that HSFA7b controls transcriptional activation of *WUS* at the SAM after HS through direct regulation of the *SYD* chromatin remodelling factor.

## Results

### *HSFA7a* and *HSFA7b* display transcriptional heat stress memory at the SAM

Transcriptome analysis of shoot apices of Arabidopsis wild-type (Col-0) seedlings^10^ revealed induced expression of *HSFA7a* and *HSFA7b* after priming HS, and hyper-induced expression in response to a more severe triggering HS, demonstrating transcriptional HS memory^6^. *HSFA7b* showed a stronger response to thermopriming than *HSFA7a* (**Fig. 1a**). To confirm the transcriptional induction of both genes, we analyzed their expression at the SAM of Col-0 plants subjected to a previously established thermomemory assay (**Supplemental Fig. 1**)^7,10^, at 0.5 h after priming and 0.5 h after triggering treatments, by qRT-PCR. This confirmed enhanced expression of *HSFA7a* and *HSFA7b* after priming HS, and strong induction after the triggering HS (**Fig. 1b, c**). Importantly, *HSFA7a/b* expression was higher in plants subjected to both priming and triggering (PT plants) than in plants exposed only to a severe triggering HS (T plants). To gain information of *HSFA7a* and *HSFA7b* expression at the SAM in higher spatial resolution, we performed RNA *in situ* hybridization (**Fig. 1d, e**). We did not detect *HSFA7a* or *HSFA7b* expression at the Col-0 SAM under control condition (C plants). However, both genes were rapidly induced at the SAM of P plants directly after the priming HS, and expression was hyper-induced at the SAM of PT plants after triggering compared to T plants, confirming that *HSFA7a* and *HSFA7b* are transcriptional HS memory genes. Our data also confirmed higher expression of *HSFA7b* than *HSFA7a* at the SAM after heat treatment. Next, we tested expression of both *HSFs* in whole seedlings, cotyledons, and shoot and root apices of PT plants at 0.5 h after triggering (**Fig. 1f, g**). Transcripts of *HSFA7a* and *HSFA7b* were upregulated in all organs, suggesting them as HS memory components throughout the whole plant. However, expression of *HSFA7a* and *HSFA7b* was most induced at the SAM, indicating this meristematic organ as their predominant localization to generate HS memory. As HSFA7a and HSFA7b share 59% amino acid sequence similarity (**Supplemental Fig. 2**), we tested their capacity for interaction by yeast two-hybrid assay. **Figure 1h** shows that both TFs interact to form a heterodimer likely involved in regulating target genes.

**Fig. 1:**
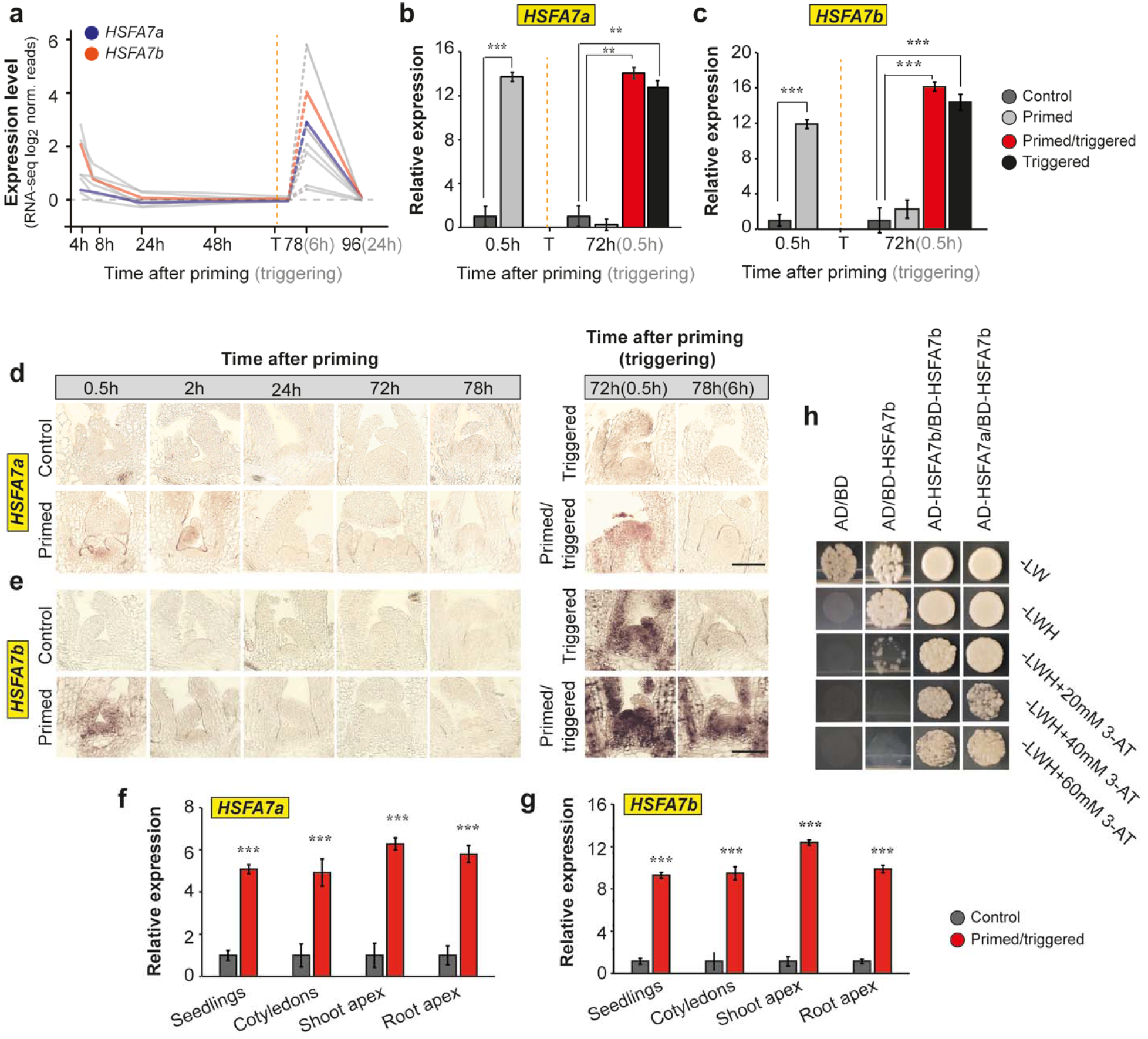
*HSFA7a* and *HSFA7b* acts as HS memory genes at the shoot apical meristem (SAM). **a**, Expression profiles of *HEAT SHOCK TRANSCRIPTION FACTORS* (*HSFs*) (grey) including *HSFA7a* (blue) and *HSFA7b* (orange) at the SAM of Col-0 plants at 4, 8, 24, 48, 78 h, and 96 h after priming (6 and 24 h after triggering) normalized to untreated, control plants. **b, c**, Expression level of *HSFA7a* **(b)** and *HSFA7b* **(c)** at the SAM of Col-0 plants after priming and triggering treatments obtained by qRT-PCR. **d, e,** RNA *in situ* hybridization using *HSFA7a* **(d)** and *HSFA7b* **(e)** as probes on longitudinal sections through meristems of control, primed, primed/triggered (PT), and triggered plants. Scale bars, 100 μm. **f, g** Tissue-specific expression of *HSFA7a* **(f)** and *HSFA7b* **(g)** in 8-day-old control and PT plants at 0.5 h after triggering treatment analyzed by qRT-PCR. **h,** Yeast-2-hybrid assay depicts the protein-protein interaction of functional HSFA7b with HSFA7b (homodimer formation) and functional HSFA7b with HSFA7a (heterodimer formation). Abbreviations: –LWH = SD-leucine, tryptophan and histidine. 3-AT = 3-amino-1,2,4-triazole. BD = GAL4 binding domain, AD = GAL4 activation domain. Error bars indicate s.d. (*n* = 3). Asterisks indicate statistically significant differences (Student *t*-test: \*\**P* ≤ 0.01 and \*\*\**P* ≤ 0.001) compared to the control conditions. In (a-c) the vertical dashed line represents the time point of triggering (T) treatment. Time is given in hours (h) after priming (black color) and triggering (grey color) treatments.

### HSFA7b maintains thermomemory

To test whether HSFA7a/b support HS memory, we subjected *hsfa7a* and *hsfa7b* single, and *hsfa7a hsfa7b* double mutants to the thermopriming assay (**Fig. 2a, Supplemental Fig. 3**). When Col-0 seedlings were subjected to a triggering HS in the absence of a prior priming stimulus, cotyledons bleached and no new leaves formed even after an extended cultivation period, as reported^7,10^. In contrast, Col-0 PT seedlings established new leaves at the SAM after the triggering HS and shoot development progressed^7,10^ (**Fig. 2a**). Importantly, while growth of *hsfa7a, hsfa7b, hsfa7a hsfa7b*, and Col-0 plants was similar in control and primed conditions, we observed impaired growth recovery, reduced survival, and decreased biomass after triggering HS in previously primed *hsfa7a hsfa7b* plants compared to Col-0, clearly demonstrating defective HS memory in the double mutant (**Fig. 2a-c**). Our conclusion was substantiated by high-resolution 3D imaging of the dynamic plant growth^16^: the PT *hsfa7a hsfa7b* mutant produced significantly smaller rosettes than Col-0 by the end of the imaging period (i.e., 11 days after triggering), concomitant with a decreased relative rosette expansion growth rate (**Fig. 2d, Supplemental Fig. 4a**). Of particular importance, however, is that both, basal and acquired thermotolerance were not altered in *hsfa7a hsfa7b* seedlings compared to wild type (**Supplemental Fig. 4b, c**), proving a specific involvement of *HSFA7a* and *HSFA7b* in thermomemory rather than a general response to HS. Furthermore, increased yellowing of PT *hsfa7a hsfa7b* leaves in response to triggering HS suggested activation of chlorophyll breakdown, senescence, and programmed cell death in these plants. Indeed, chlorophyll content was significantly lower in PT *hsfa7b* single and *hsfa7a hsfa7b* double mutants compared to Col-0 seven days after the triggering HS (**Fig. 2e**). We also tested whether thermopriming induces cell death 24 h after triggering HS by Trypan Blue staining of cotyledons (**Supplemental Fig. 5a**). While only 18% of Col-0 PT plants displayed lesions, 42%, 43%, and 75% of *hsfa7a* and *hsfa7b* single and *hsfa7a hsfa7b* double mutants, respectively, showed dead cells (**Supplemental Fig. 5b**), demonstrating that thermopriming triggers cell death in the absence of HSFA7a/b. To analyze whether lack of functional HSFA7a and/or HSFA7b affects thermopriming capacity of the SAM, we analyzed expression of the HS memory marker *HEAT SHOCK PROTEIN 17.8 (HSP17.8)^10^* by RNA *in situ* hybridization (**Fig. 2f**). Expression of *HSP18.2* was similar at the SAM of Col-0, *hsfa7a, hsfa7b*, and *hsfa7a hsfa7b* plants after priming HS. In contrast, after triggering, SAM-located *HSP18.2* expression was reduced in both *hsfa7* single-gene mutants and virtually absent in the *hsfa7a hsfa7b* double mutant, demonstrating the importance of both HSFs for establishing thermomemory at the SAM.

**Fig. 2:**
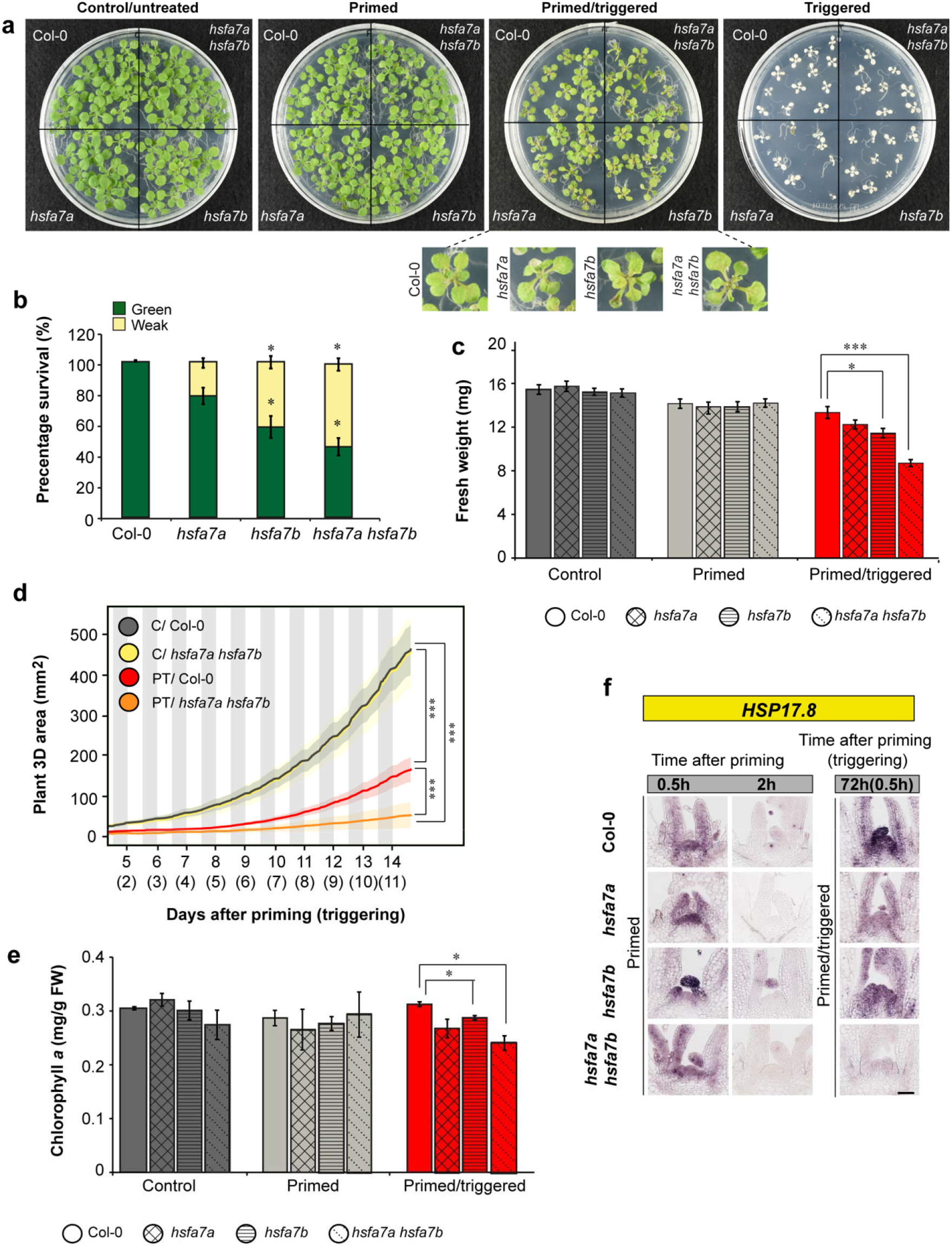
HSFA7a and HSFA7b are required for thermomemory at the SAM. **a,** Growth recovery phenotypes of the control (C), primed (P), primer/triggered (PT), and triggered (T) Col-0 wild-type and the *hsfa7a, hsfa7b*, and *hsfa7a hsfa7b* mutant plants. Note, no new leaf formation was observed in the thermoprimed *hsfa7a hsfa7b* double mutant (zoomed images in the panel). Pictures were taken at 7 days after triggering (DAT, 10 days after priming). **b,** Percentage survival of seedlings in different phenotype classes analyzed at 7 DAT. ‘Green’ represents seedlings in which shoot regeneration continued and almost the entire plant was green; ‘weak’ represents seedlings in which shoot regeneration was weak and plants were mostly pale; Error bars represent means s.d. (*n* = 3). **c,** Fresh weight of C, P, and PT Col-0, *hsfa7a, hsfa7b*, and *hsfa7a hsfa7b* mutant plants. Measurements were taken 7 DAT. Error bars indicate means s.d. (*n* = 3). **d,** 3D total plant area of C and PT Col-0 and *hsfa7a hsfa7b* double mutant plants measured over time (*n* ≥ 6 for each condition). Lines and color-shaded areas represent mean and s.d., respectively. **e,** Chlorophyll *a* content analyzed in C, P, and PT Col-0 and mutant plants at 10 days after triggering. Error bars indicate s.d. (*n* = 3). **f,** RNA *in situ* hybridization using *HEAT SHOCK PROTEIN 17.8* (*HSP17.8*) as probe on longitudinal sections through meristems of P and PT Col-0, *hsfa7a, hsfa7b*, and *hsfa7a hsfa7b* mutant plants. Asterisks indicate statistically significant difference (Student’s *t*-test: \**P* ≤ 0.05; \*\*\**P* ≤ 0.001) compared to Col-0 under the same conditions.

### Transcriptome analysis of *hsfa7a* and *hsfa7b* shoot apices

To determine the molecular basis of the phenotypic changes in the *hsfa7a/b* mutants, we performed RNA-sequencing (RNA-seq) of shoot apices 0.5 h after priming HS and 0.5 h after triggering HS (**Supplemental Fig. 6; Supplemental Data 1**). In response to priming and triggering, several hundred genes were significantly differentially expressed between *hsfa7a/b* mutants and Col-0. To identify high-confidence genes, we introduced a fold change criterion in gene expression of |FC|> 1.5) (**Supplemental Table 1**). Moreover, as *hsfa7a/b* mutants are impaired in HS memory but not in basal or acquired thermotolerance, we focused on the differentially expressed genes (DEGs) after triggering HS (**Fig. 3a**). Firstly, we generated a Venn diagram of the DEGs identified in *hsfa7a, hsfa7b*, and *hsfa7a hsfa7b* (**Fig. 3b**). The intersection of differentially expressed transcripts shared between all three mutants after triggering treatment included only 12 genes. In particular, 28 DEGs (of 63) were specific for *hsfa7a*, 20 (of 53) for *hsfa7b*, and 202 (of 252) for *hsfa7a hsfa7b* after triggering HS. Next, we intended to identify DEGs specific for the PT *hsfa7a hsfa7b* double mutant (**Supplemental Data 2**). Thus, we generated a heat map of the 202 genes (**Fig. 3c**) only affected in *hsfa7a hsfa7b* shoot apices after triggering HS. Evidently, the expression profiles of *hsfa7a hsfa7b-specific* genes strongly differ from those of Col-0 and *hsfa7a* and *hsfa7b* single mutants, suggesting that the HSFA7a – HSFA7b complex controls a different set of genes in response to HS than the individual HSFA7a/b proteins. Genes like *ETHYLENE RESPONSIVE ELEMENT BINDING FACTOR 1A (ERF1A), ERF2*, and *ERF104* are included in the ‘ethylene-activated signalling pathway’ category, indicating that the response to ethylene is affected by thermopriming at the SAM.

**Fig. 3:**
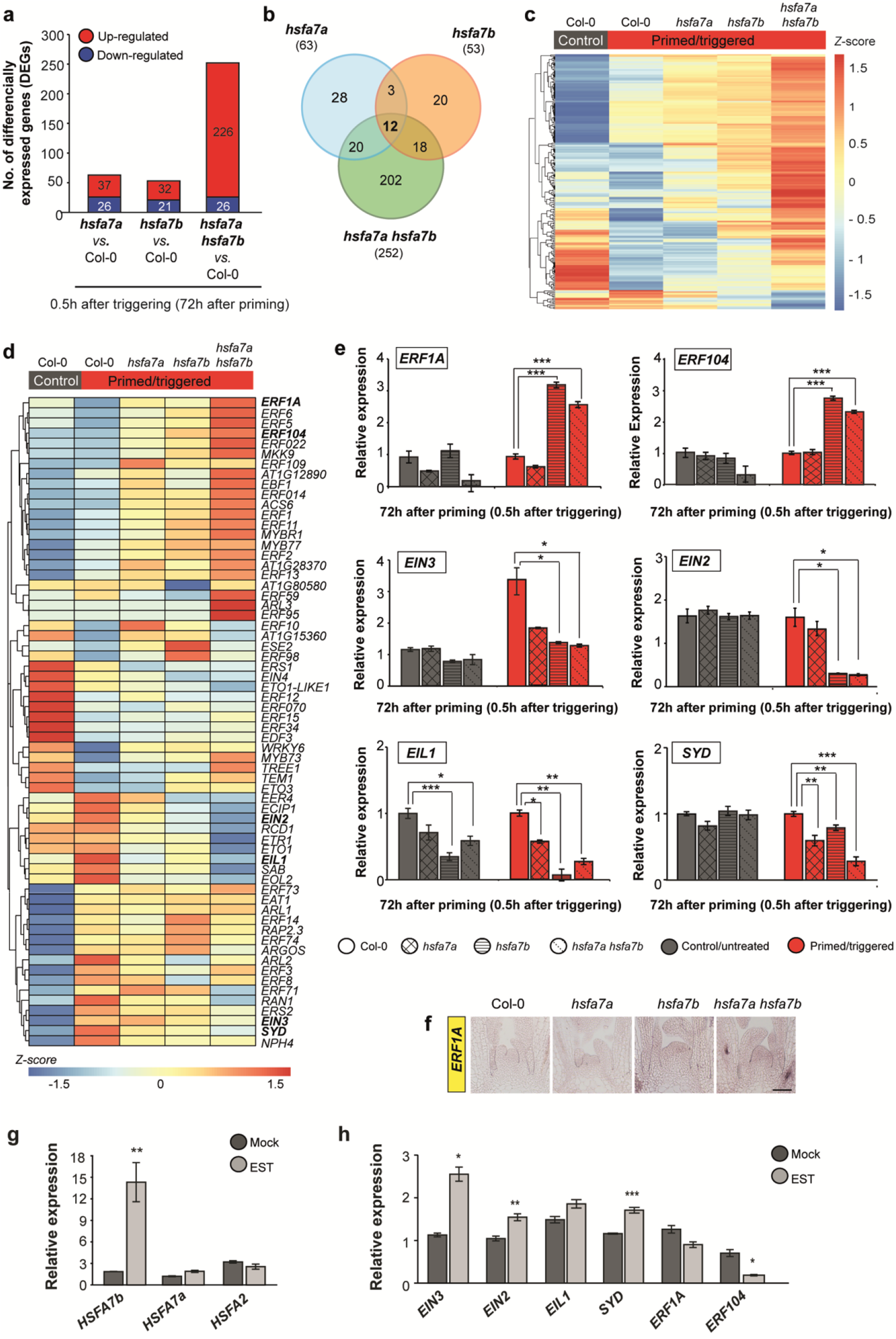
Thermopriming affects expression of ethylene response genes at the shoot apical meristem in an HSFA7b-dependent manner. **a,** Total number of differentially expressed genes (DEGs; FDR < 0.05 and |FC| > 1.5) at the shoot apices of primed and primed/triggered (PT) *hsfa7a, hsfa7b*, and *hsfa7a hsfa7b* mutants compared to PT Col-0 wild-type plants with number of up-(red) and down-regulated (blue) genes. **b,** Venn diagrams of overlapping DEGs at 0.5 h after triggering in *hsfa7a/b*, and *hsfa7a hsfa7b* mutants compared to control Col-0 plants. **c,** Heat map of 202 DEGs identified to be specific for PT *hsfa7a hsfa7b* double mutant and their relative expression (*Z*-score normalized). **d,** Heat map depicting relative expression (*Z*-score normalized) of genes of the GO term category ‘ethylene biosynthesis and response’ in the Col-0 control and PT Col-0, *hsfa7a, hsfa7b*, and *hsfa7a hsfa7b* shoot apices. **e,** Expression level of selected ethylene-related genes analysed by qRT-PCR at the SAM of control and PT Col-0, *hsfa7a/b*, and *hsfa7a hsfa7b* plants at 0.5 h after triggering. **f,** RNA *in situ* hybridization using *ETHYLENE RESPONSIVE FACTOR 1A* (*ERF1A*)-specific probe at longitudinal sections through meristems of PT Col-0, *hsfa7a/b*, and *hsfa7a hsfa7b* plants. Scale bar, 100 μm. **g, h,** Expression level of (g) *HEAT SHOCK TRANSCRIPTION FACTOR A7a (HSFA7a), HSFA7b, HSFA2*, and (h) ethylene-related genes analysed in *HSFA7a-IOE* and mock-treated plants. Error bars represent ± s.d. (*n* = 3). Asterisks indicate statistically significant difference (Student’s *t*-test, *P* < 0.05, \*\**P* < 0.01, and \*\*\**P* < 0.001) compared to Col-0 under the same condition (b) and compared to mock treatment (d and e).

### Thermopriming affects expression of ethylene-related genes in an HSFA7b-dependent manner

The finding that expression of ethylene-related genes is affected by triggering HS in PT *hsfa7a hsfa7b* plants prompted us to investigate whether ethylene signalling plays a role in HS memory at the SAM. **Figure 3d** shows that expression of ethylene signalling genes clearly differed between *hsfa7a/b* single and double mutants and Col-0. The most pronounced changes were observed in *hsfa7b* and *hsfa7a hsfa7b*, suggesting that expression of ethylene-related genes mostly requires HSFA7b, with a smaller impact by HSFA7a. We confirmed this by testing the expression of *ERF1A, ERF104, ERF11, ETHYLENE-INSENSITIVE 3* and *2 (EIN3* and *EIN2), ETHYLENE-INSENSITIVE3-LIKE1 (EIL1*), and *SYD* in meristems of C and PT plants at 0.5 h after triggering (**Fig. 3e, Supplemental Fig. 7a**). Of note, expression of *ERF1A, ERF11*, and *ERF104* was elevated at the SAM of PT *hsfa7b* and *hsfa7a hsfa7b* plants compared to untreated and PT Col-0 plants. Furthermore, we observed reduced expression of *EIN3, EIN2, EIL1*, and *SYD* at the SAM of *hsfa7b* and *hsfa7a hsfa7b* compared to PT Col-0 (**Fig. 3e**). Downregulation of these genes after triggering HS was also observed in *hsfa7a*, however the response was much weaker to that observed in *hsfa7b* single and double mutants. We additionally performed RNA *in situ* hybridization for *ERF1A* (**Fig. 3f**). After a triggering HS, *ERF1A* was weakly expressed at the SAM of PT Col-0 plants while its expression greatly increased in *hsfa7b* and *hsfa7a hsfa7b* mutants, compared to Col-0 and *hsfa7a*. The results support the model that HSFA7b controls the expression of ethylene-related genes at the SAM during thermopriming. To validate this, we generated a β-estradiol (EST)-inducible *HSFA7b* overexpression line (*HSFA7b-IOE;***Supplemental Fig. 7b**) and performed RNA-seq after EST induction which yielded 1,219 up- and 548 downregulated genes, respectively, compared to mock treatment (**Supplemental Fig. 7c**). The clustered heat map of ethylene-related genes revealed their altered expression of EST-induced *HSFA7b-IOE* seedlings (**Supplemental Fig. 7d**). The expressional changes of *HSFA7b, ERF1A, ERF104, EIN3, EIN2, EIL1*, and *SYD* in *HSFA7b-IOE* seedlings were confirmed by qRT-PCR (**Fig. 4g, h, Supplemental Fig. 7e**). Collectively, our data suggest that HSFA7b, with a minor impact from HSFA7a, regulates expression of ethylene signalling-related genes at the SAM during thermomemory. We, therefore, tested whether changes in ethylene level alter molecular and/or morphological phenotypes of *hsfa7b* and *hsfa7a hsfa7b* mutants. To induce ethylene response, we grew Col-0*, hsfa7a, hsfa7b*, and *hsfa7a hsfa7b* seedlings in the presence or absence of the ethylene precursor 1-aminocyclopropane-1-carboxylic acid (ACC) (**Fig. 4a**). Without ACC, all plants grew normally, and with ACC, root growth was reduced in all plants in a dose-dependent manner (**Fig. 4a, b**). All mutants showed a stronger response to ACC treatment than Col-0, and the highest effect was found in *hsfa7a hsfa7b*, suggesting that a lack of HSFA7a/b increases the sensitivity of plants towards ethylene (**Fig. 4a, b**). In addition, expression of *HSFA7b*, but not *HSFA7a*, was induced at the shoot apex in response to ACC treatment (**Supplemental Fig. 7f**). Collectively, our data demonstrate that HSFA7b is more relevant than HSFA7a for thermomemory of the SAM.

**Fig. 4:**
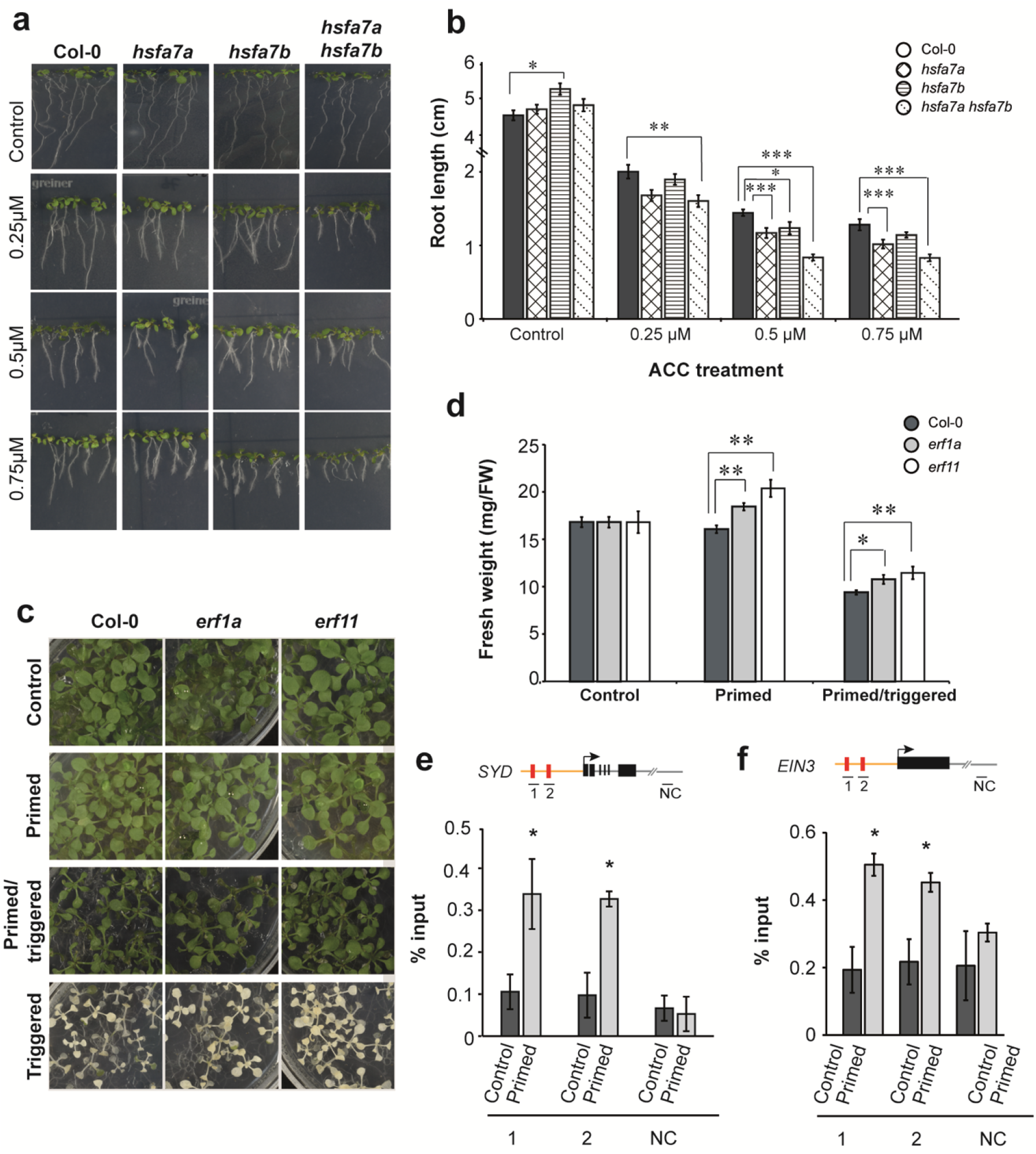
HSFA7b controls ethylene response at the SAM during thermomemory. **a,** Col-0, *hsfa7a, hsfa7b* single and *hsfa7a hsfa7b* double mutant seedlings grown on 0.5 MS medium supplemented with or without 0.25, 0.5, and 0.75 μM 1-aminocyclopropane-1-carboxylic acid (ACC). Images were taken seven days after germination. **b,** Root length of Col-0 and mutant plants grown on increasing concentration of ACC. **c,** Phenotype of control (C), primed (P), primed/triggered (PT), and triggered (T) Col-0, *erf1a*, and *erf11* mutant plants. Images were taken 10 days after triggering. **d,** Fresh weight of C, P, and PT Col-0, *erf1a*, and *erf11* mutants. **e, f,** Binding of HSFA7b to the promoter regions of **(e)** *SPLAYED* (*SYD*) and **(f)** *ETHYLENE INSENSITIVE 3* (*EIN3*) under primed conditions compared to control, determined by ChIP-qPCR. Y-axis represents the relative enrichment compared to the input (in %) and X-axis represents the genomic regions depicted in the schematics. The schematic representation of each gene depicts the regions analyzed by ChIP-qPCR (arrow, TSS; yellow line, promoter; black boxes, exon; red boxes, heat shock elements; grey line, 3’ UTR; NC; negative control). Error bars indicate s.d. (*n* = 3). Asterisks indicate statistically significant difference compared to control (Student’s *t*-test: \**P* ≤ 0.05 and \*\**P* < 0.01).

To unravel whether changes in ethylene response affect thermomemory, we subjected the ethylene-signalling and response mutants *erf1a* and *erf11* to the thermomemory assay (**Fig. 4c**). Both mutants exhibited a better recovery after priming plus triggering HS (PT) than Col-0 (**Fig. 4c, d**).

To confirm the transcriptional regulation of ethylene-related genes by HSFA7b, we generated a *pHSF7b:HSFA7b:GFP* transgenic line (**Supplemental Fig. 8**) and performed chromatin immunoprecipitation - quantitative polymerase chain reaction (ChIP-qPCR) analysis on selected target genes (*EIN3* and *SYD*) at 0.5 h after priming HS and in control seedlings (**Fig. 4e, f**). Binding of HSFA7b was highly enriched in *EIN3* and *SYD* promoters, demonstrating direct regulation of both genes, and thus the ethylene response, during thermopriming.

### HSFA7b controls meristem maintenance during thermopriming

Our results revealed that HSFA7b directly binds the promoter of the chromatin remodelling factor *SYD* to activate its expression. Since *SYD* controls *WUS* expression through direct promoter binding^15^, we investigated whether HSFA7b might activate *WUS* by regulating *SYD* expression. In Col-0 plants, *WUS* expression (qRT-PCR) was significantly higher in P and PT than control plants, demonstrating that thermopriming activates *WUS* transcription at the SAM (**Fig. 5a**). RNA *in situ* hybridization confirmed transcriptional activation of *WUS* (**Fig. 5b**). Interestingly, we observed transient ectopic expression of *WUS* at the SAM of P and PT plants, demonstrating that it is not restricted to the organizing center anymore after HS (**Fig. 5b**). Importantly, *WUS* transcript was absent from seedlings subjected to only the triggering HS, demonstrating that priming HS protects the SAM from the terminal effects of an otherwise lethal HS. As we previously demonstrated, the SAM with its stem cells collapses directly after triggering HS in the absence of a prior priming^10^, our data suggest that temporal upregulation of *WUS* in response to thermopriming is essential for plant survival. Next, we tested whether activation of *WUS* at the SAM in response to thermopriming requires HSFA7b. qRT-PCR analysis on dissected shoot apices showed a strong downregulation of *WUS* transcript in PT *hsfa7a hsfa7b* double mutants compared to PT Col-0 plants, while *WUS* expression was unchanged in control/untreated *hsfa7a hsfa7b* plants (**Fig. 5c**). Furthermore, *SHOOTMERISTEMLESS* (STM) regulates SAM maintenance independently of the SYD-WUS pathway^15^; we, therefore, analysed *STM* expression by qRT-PCR and RNA *in situ* hybridization at the SAM of Col-0 and *hsfa7* mutants (**Fig. 5d-f**). Expression of *STM* was greatly reduced in P and PT plants compared to control, and undetectable in the SAM of only-triggered plants (**Fig. 5d, e**). Importantly, *STM* expression was similar in PT Col-0 and all *hsfa7* mutant plants (**Fig. 5f**), suggesting that its decreased expression at the SAM of P and PT plants does not require functional HSFA7b. Taken together, our data suggest that HSFA7b controls meristem maintenance during HS memory *via* the SYD-WUS pathway.

**Fig. 5:**
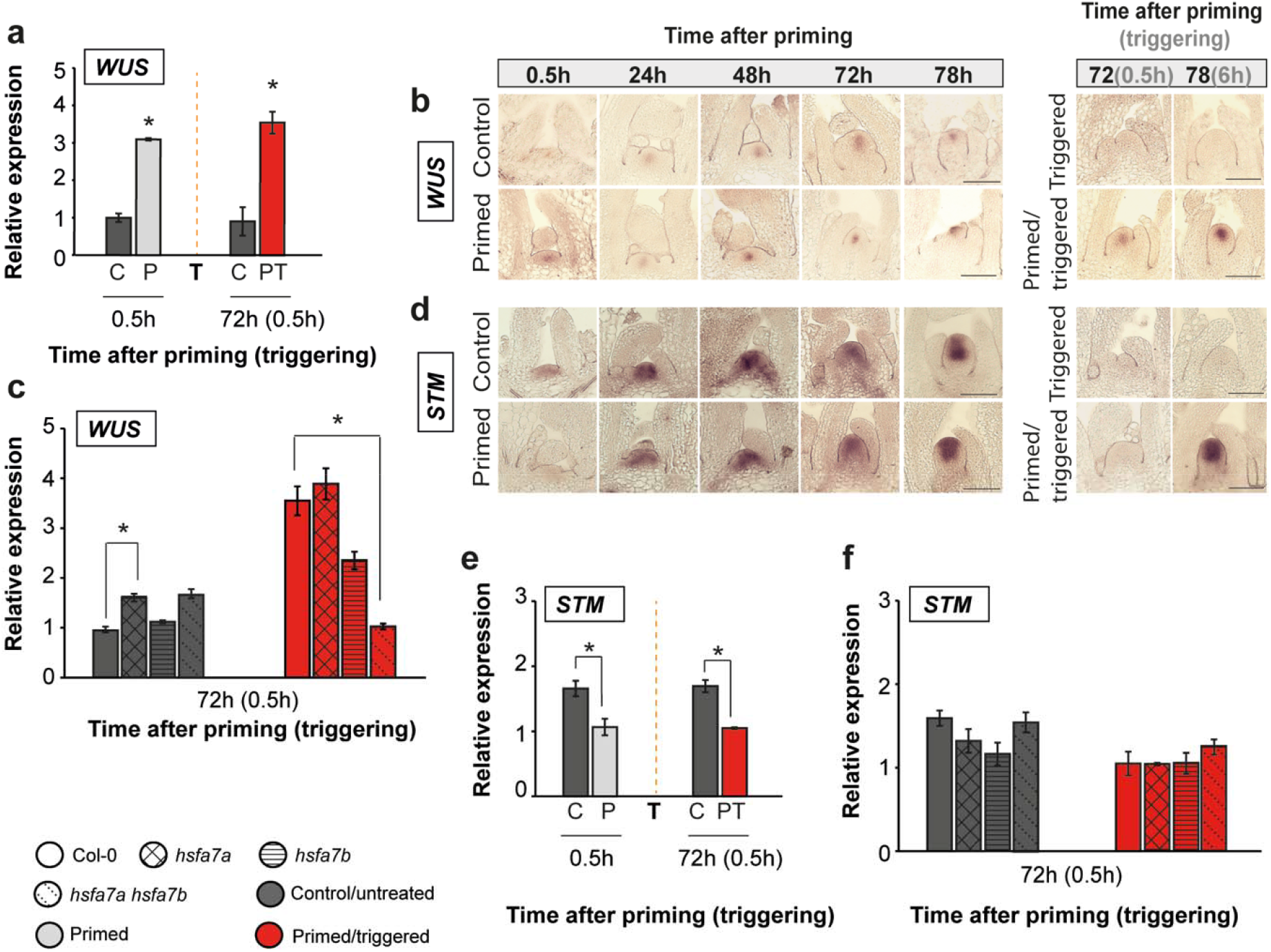
HSFA7b regulates *WUSCHEL* (*WUS*) expression at the shoot apical meristem (SAM) through direct transcriptional activation of *SPLAYED* chromatin remodelling factor. **a,** Expression level of *WUS* at the SAM of control (C), primed (P), and primed/triggered (PT) Col-0 wild-type plants. **b,** RNA *in situ* hybridization using a *WUS*-specific probe on longitudinal sections trough meristems of C, P, PT, and triggered plants. Note that the *WUS* transcript is absent at the SAM of plants exposed only to triggering (T) treatment. **c,** Expression of *WUS* at the SAM of C and PT Col-0, *hsfa7a, hsfa7b*, and *hsfa7a hsfa7b* mutant plants at 0.5 h after triggering. **d,** RNA *in situ* hybridization using a *SHOOTMERISTEMLESS* (*STM*)-*specific* probe on longitudinal sections trough meristems of C, P, PT, and triggered plants. **e,** Expression level of *STM* at the SAM of C, P, and PT Col-0 plants at 0.5 h after priming and 0.5 h after triggering treatments. **f,** Expression level of *STM* at the SAM of C and PT Col-0, *hsfa7a, hsfa7b*,and *hsfa7a hsfa7b* mutant plants at 0.5 h after priming and 0.5 h after triggering treatments. Scale bar, 100 μm (b, e). Error bars represent ± SEM; (*n* = 3). Asterisks indicate statistically significant difference (Student’s *t*-test, \**P* ≤ 0.05 and \*\**P* < 0.01) compared to control under same conditions.

**Fig. 6:**
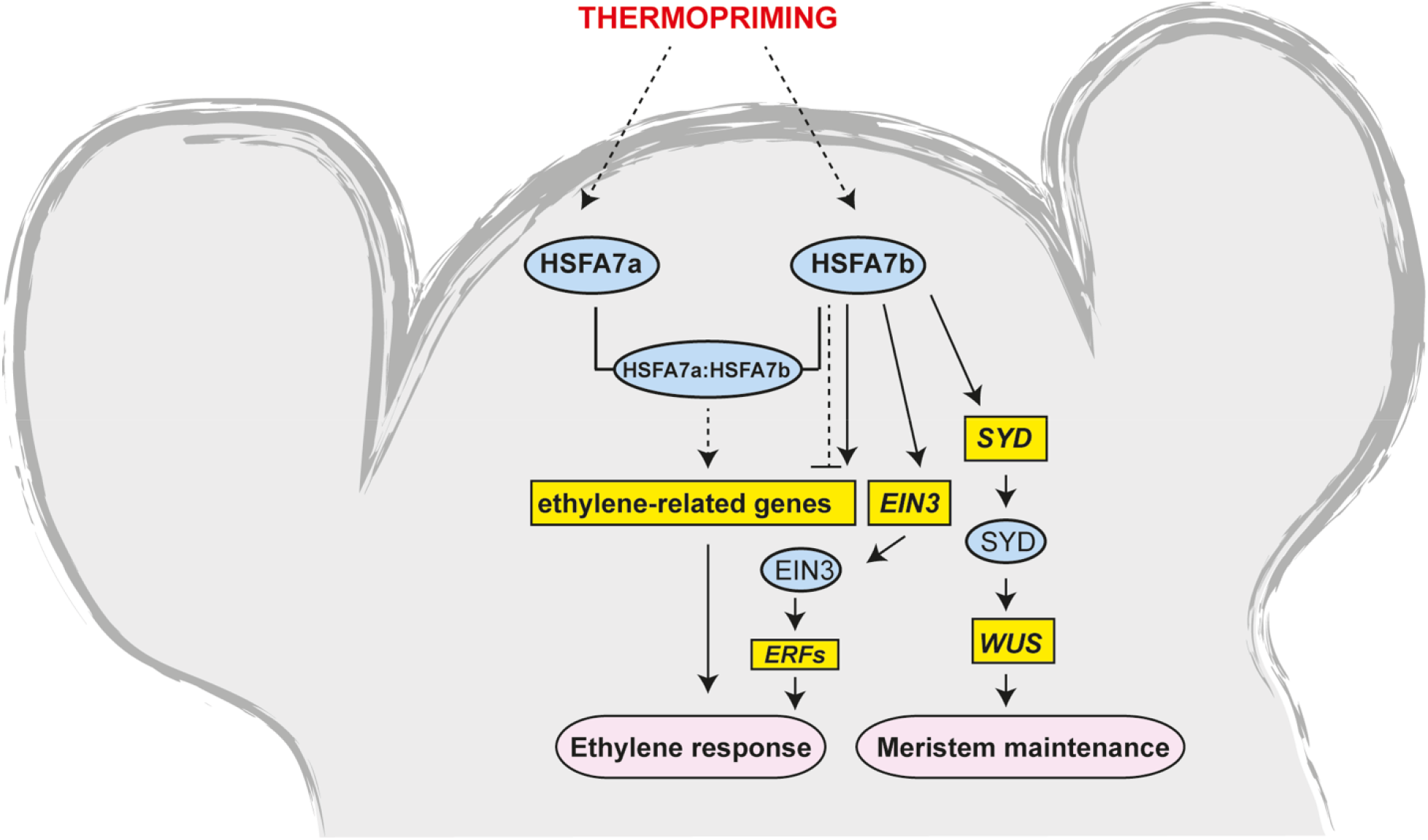
Schematic model showing the regulation of the ethylene response by HSFA7b at the shoot apex during thermomemory. HSFA7b controls ethylene response by binding to HSEs in the promoter regions of ethylene-related genes including *ETHYLENE-INSENSITIVE 3* (*EIN3*) and *SPLAYED* (*SYD*) to activate a transcriptional HS cascade and cover thermotolerance at the SAM. The direct transcriptional activation of *SYD* by HSFA7b leads to upregulation of *WUSCHEL (WUS*) transcription at the SAM. Yellow rectangular boxes indicate genes, blue ovals indicate proteins, and pink rectangles with rounded ends indicate cellular processes. Solid lines, direct interactions; dashed lines, indirect interactions.

## Discussion

The global temperature increase poses serious threat to agriculture. Recent research suggests that climate change triggered mismatches in above- and below-ground plant phenology and that shoots and roots respond differently to the environmental input^17^. It has, therefore, become imperative to study the tissue-specific responses to HS to improve the thermotolerance of crops.

We show that SAM-expressed *HSFA7a* and *HSFA7b* generate HS memory in Arabidopsis seedlings: *hsfa7b* and *hsfa7a hsfa7b* mutants have reduced thermomemory compared to wildtype plants, while basal and acquired thermotolerance are not affected. The HS memory of the *hsfa7a hsfa7b* SAM is strongly impaired as e.g. evidenced by reduced expression of the memory gene *HSP18.2* after triggering HS. The main role of molecular chaperone HSPs is to protect plants from devastating effects imposed by the stressor (here, HS)^18^.

An important result of our transcriptome analysis is that HSFA7a and HSFA7b control different sets of genes in response to thermopriming indicating dynamic but as yet vaguely understood events acting at the SAM during thermomemory, or abiotic stress responses in general. A key finding is that HSFA7b controls ethylene response at the SAM during thermopriming, while HSFA7a has only a minor impact on the expression of ethylene-related genes. For example, ethylene response genes *ERF1A* and *ERF11* are strongly upregulated only at the SAM of *hsfa7b* and *hsfa7a hsfa7b* mutants after triggering HS. *ERF11* negatively regulates abiotic stress tolerance in plants^19,20^. Accordingly, *erf11* seedlings display better growth recovery than Col-0 after priming/triggering HS.

Direct regulation of the ethylene response at the SAM by HSFA7b is supported by ChIP-qPCR which revealed direct interaction with key players of ethylene signalling, *EIN3* and *SYD*. EIN3, the key transcriptional regulator of ethylene signaling, binds to the promoters of *ERF95* and *ERF97* to mediate thermotolerance^21^. Thus, HSFA7b appears to control HS memory by affecting ethylene signalling by binding to the *EIN3* promoter to activate an ERF transcriptional cascade at the SAM. Furthermore, our data might suggest that ethylene production is increased in *hsfa7b* and *hsfa7a hsfa7b* mutants after PT treatment as concluded from the hypersensitivity of the *hsfa7a hsfa7b* double mutant in response to ACC treatment. Previous studies reported an accumulation of ethylene after HS^22^–^24^. An increased production of ethylene under HS correlates with an acceleration of leaf senescence^25^. In accordance with this, we found that chlorophyll breakdown, the activation of senescence, and programmed cell death occur more rapidly in the double mutant than wild type after the triggering HS.

Our study shows that HSFA7b contributes to meristem maintenance during HS by directly binding to HSEs in the *SYD* promoter. As a chromatin remodelling factor, SYD regulates the pool of stem cells at the shoot apex by transcriptionally activating *WUS*^15^. In accordance with this, expression of both *SYD* and *WUS* was strongly downregulated at the SAM of PT *hsfa7a hsfa7b* mutant seedlings. Thus, the upregulation of *WUS* expression in wild-type plants after priming and triggering treatments suggests a key role of WUS in the SAM’s thermoprotective processes. The elevated expression of *WUS* in PT Col-0 plants is consistent with the downregulation of *CLV1* and *CLV3* at the SAM in response to thermopriming and their inhibitory role on *WUS* transcription^10,15^. As expression of *STM* remained unchanged at the SAM of the *hsfa7b* mutant, HSFA7b appears to specifically act *via* the WUS pathway during HS. Interestingly, ectopic *WUS* expression also occurs in *bru1*, a mutant lacking another chromatin remodelling factor, BRUSHY1, and *bru1* is impaired in HS memory^26^. Similarly, FORGETTER 1 mediates HS-induced chromatin memory by interacting with the CHROMATIN REMODELING FACTORs 11 and 17 as well as with BRAHMA (BRM)^26^. The findings suggest that chromatin remodelling factors play an eminent role in meristem maintenance and HS memory^27^. In this regard it is important to decipher how upstream regulators like e.g. transcription factors control the expression of chromatin remodelling genes. One of them is HSFA2 which directly regulates expression of *BRM* during thermomemory^3^. Our study shows that HSFA7b directly controls expression of *SYD* during HS memory, thus demonstrating that it act as an important TF to directly link HS signals with meristem maintenance.

## Acknowledgements

B.M.-R. thanks the Deutsche Forschungsgemeinschaft (DFG) for funding Collaborative Research Centre 973 ‘Priming and Memory of Organismic Responses to Stress’ (www.sfb973.de) and the European Union’s Horizon 2020 Research and Innovation Programme for funding PlantaSYST (SGA-CSA No. 739582; FPA No. 664620). B.M.-R. thanks the Max Planck Institute of Molecular Plant Physiology and the University of Potsdam for financial support. B.M.-R. and S.J. thank the International Max Planck Research School ‘Primary Metabolism and Plant Growth’ (IMPRS-PMPG) for support. J.J.O. thanks the DFG for funding (OL 767/1-1). We thank Eike Kamann for cloning and technical assistance, and Sarah Richard and Philip Cieslak for general lab work.

## Author contributions

J.J.O. and B.M.-R. conceived the study and designed the experiments. S.J. carried out experiments, except some RNA *in situ* hybridizations (J.J.O). F.A. performed RNA-seq analysis with contributions from A.K. F.A. and D.B. measured plant 3D growth. M.G.A. measured chlorophyll content. F.F. generated the *pHSFA7b::HSFA7b:GFP* transgenic line. J.J.O. wrote the manuscript with contributions from S.J. and B.M.-R. All authors read and commented on the manuscript before submission.

## Material and Methods

### Plant material and growing conditions

*Arabidopsis thaliana* Col-0 seedlings were grown on 0.5 Murashige and Skoog (MS) medium with 1% sucrose under long-day (LD, 16 h light/8 h dark) conditions at 22°C with photosynthetically active radiation at 160 μmol m^-2^ s^-1^ in a controlled growth chamber (Fitotron SCG 120, Weiss Technik, Loughborough, U.K.). Thermomemory assay, basal thermotolerance, and acquisition of thermotolerance were performed as previously described^9,10^. Briefly, five-day-old seedlings were subjected to the priming treatment (1.5 h at 37°C, 1.5 h at 22°C, 45 min at 44°C) at 6 h after dawn, followed by a recovery/memory phase at 22°C for three days, and triggering stimulus (1.5 h at 44°C) at 9 h after dawn. The *hsfa7a* (SALK_080138) and *hsfa7b*(SALK_152004) mutants were previously reported^28^ and were obtained from the Nottingham Arabidopsis Stock Centre collection. Homozygous lines were confirmed by PCR and the primer sequences for genotyping are listed in **Supplemental Table 2**. The *hsfa7a hsfa7b* double mutant was generated by crossing the homozygous *hsfa7a* and *hsfa7b* single mutants. For generating the *HSFA7b* complementation line (*pHSFA7b:HSFA7b-GFP*), a modified pGreen0229 plant transformation vector containing a C-terminal GFP tag was used^29^. The *HSFA7b* open reading frame (without the stop codon) and its 2-kb promoter region were amplified by PCR from Arabidopsis Col-0 leaf genomic DNA. For generating the β-estradiol (EST)-inducible *HSFA7b* overexpression line (*HSFA7b-IOE*), the pER8 vector was used^30^.

### Determination of plant size, RER, and fresh weight

Plants were classified into two phenotype classes based on their recovery phenotype at seven days after priming/triggering treatment. The seedlings in which shoot regeneration continued and the shoot remained green were classified as ‘green’, while seedlings with weak shoot regeneration and a mostly pale appearance were classified as ‘weak’. Fresh weight of rosettes of control (C), primed (P), and primed/triggered (PT) Col-0 and mutant plants was measured seven days after the triggering treatment.

Plant 3D rosette area and relative expansion growth rate (RER) of Col-0 wild type (*n* = 10 for C; *n*= 6 for PT) and *hsfa7a hsfa7b* mutant plants (*n* = 12 for C; *n* = 6 for PT) were analyzed using an established 3D camera-based imaging system^16^. Plants were transferred to soil one day after triggering and continuously imaged in a growth chamber (model E-36L; Percival Scientific) for several days as previously described^10^.

### Chlorophyll measurement

For chlorophyll measurement, whole rosettes were harvested and analyzed as previously described^31^. Assays were performed in 96-well plates and absorbances at 645 and 665 nm were determined using a Synergy microplate reader (Bio-Tek). For all assays, two technical replicates were determined per biological replicate.

### RNA *in situ* hybridization

For RNA *in situ* hybridization experiments, meristems of plants grown in LD condition were harvested in freshly prepared formaldehyde: acetic acid (FAA) fixative solution after different time points of the thermomemory assay. The samples were then transferred into embedding cassettes and fixed overnight using an automated tissue processor (Leica ASP200S, Wetzlar, Germany). After fixation, the samples were embedded in paraffin wax using the embedding system (HistoCore Arcadia, Leica, Wetzlar, Deutschland). Longitudinal sections through the apices (8 μm thickness) were prepared using a rotary microtome (Leica RM2255). Slides were stored until use for RNA *in situ* hybridization.

RNA *in situ* hybridization was done as described^32^. In order to compare the changes in gene expression, slides containing longitudinal sections through meristems of Col-0 and mutant plants were processed at the same time to exclude the possibility of variations in length of time given for probe hybridization and signal development before imaging.

### Quantitative reverse transcription PCR (qRT-PCR) and RNA-seq

Shoot and root apices, cotyledons and whole rosettes of Arabidopsis plants were collected during and after thermopriming treatment to study gene expression. Total RNA was isolated using the Trizol (Ambion/Life Technologies, Darmstadt, Germany) method. DNA digestion and cDNA synthesis were performed using Turbo DNA-free DNase I kit (Ambion/Life Technologies, Darmstadt, Germany) and the RevertAid H minus reverse transcriptase kit (ThermoFisher Scientific, Darmstadt, Germany), respectively. The qRT-PCR measurements were performed using the SYBR Green-PCR Master Mix (Applied Biosystems/Life Technologies, Darmstadt, Germany). Relative expression of each gene was analysed using a comparative cycle threshold (CT) method^33^ with a *TUBULIN2 (TUB2*) as reference gene. Primer sequences are listed in **Supplemental Table 2**.

For RNA-seq, shoot apices of Col-0, *hsfa7a, hsfa7b*, and *hsfa7a hsfa7b* were dissected in three biological replicates at 0.5 h after the priming and 0.5 h after triggering treatments. Additionally, mock and EST-treated *HSFA7b-IOE* seedlings were harvested for transcriptome analysis (n = 3). Total RNA was isolated using the *mirVana* RNA isolation kit (Ambion/Life Technologies, Darmstadt, Germany) according to the manufacturer’s protocol. Library preparation and sequencing to generate paired-end reads (2 × 100 bp) was performed by BGI Tech Solutions Co., Ltd (Hong Kong, China).

STAR program (version 2.5.2b)^34^ was used to align the reads to the Arabidopsis reference genome (*Arabidopsis thaliana*, TAIR10). The aligned reads were quantified using the HTSeq (version 0.9.1)^35^. For details of the libraries, read numbers, and alignments see Supplemental Data 1. Genome annotations were used from Araport11, RepTAS, and miRBase^36–38^. Differential gene expression analysis was done using DESeq2 (1.20.0)^39^ with criteria of FDR < 0.05 and |FC| > 1.5 for DEGs. The detailed pairwise comparisons for DEGs are listed in **Supplemental Data 2**. Heat maps plots were generated with normalized expression values generated by applying variance stabilizing transformation (VST) using DESeq2. Analysis of Gene Ontology (GO) was performed using PANTHER 16.0 - Gene list analysis http://pantherdb.org/. Genes involved in ethylene response were selected manually from The Arabidopsis Information Resource website (TAIR).

### Trypan Blue staining

Trypan Blue staining was performed 24 h after triggering treatment as previously reported^40^. Briefly, seedlings of Col-0 and mutant plants were incubated in 0.4 mg/ml Trypan Blue, dissolved in phenol/glycerol/lactic acid/water/ethanol (1:2:1:1:8) at room temperature for 30 min, and afterwards de-stained by washing three times in 90% EtOH.

### Yeast two-hybrid assay

For bait construction, the coding sequence of HSFA7a and HSFA7b were cloned in the pDEST32 vector (containing the GAL4 binding domain) using GATEWAY cloning. The positive clones were transformed in the yeast strain pJ684α. For the prey construction, the coding sequence of the functional HSFA7b was cloned in the pDEST22 vector (containing the GAL4 activation domain) using GATEWAY cloning. The positive clone was transformed in the yeast strain YM4271a. The prey with the HSFA7a protein was used from a TF library containing approximately 1,200 Arabidopsis TFs, established in vector pDEST222 in yeast strain YM4271a. For testing interactions, the yeast cells were mated in the following combinations: BD-HSFA7b - AD-HSFA7b and BD-HSFA7b - AD-HSFA7a. The empty GAL4 BD bait vector (pDEST32) and GAL4 AD prey vector (pDEST22) combination was used as a negative control. After mating for three days, the cells were plated on the following selection media: SD-Leu-Trp (mating control) and SD-Leu-Trp-His + 3AT (3-amino-1,2,4-triazole) to check for positive interactions where *HIS3* is the reporter gene and 3AT is a competitive inhibitor of the *HIS3*-encoded histidine biosynthesis enzyme.

### ChIP-qPCR

Five-day-old control and primed *pHSFA7b:HSFA7b-GFP* seedlings were used for ChIP-qPCR experiments. Briefly, ChIP-qPCR was performed as previously reported^10^. One g fresh weight of seedlings was harvested within 0.5 h after the priming treatment. The crosslinking was performed by vacuum infiltration (~ –950 millibars) for 20 min and ChIP-qPCR were performed using Diagenode’s Universal Plant kit (Diagenode, Seraing, Belgium). The shared chromatin was immunoprecipitated using anti-GFP antibody (Abcam 290), and no-antibody as a control. Three biological replicates were used for each ChIP reaction. The DNA library preparation was performed using the NEBNext^®^ Ultra^™^ II DNA Library Prep Kit for Illumina (# E7645S/L, New England BioLabs, Frankfurt am Main, Germany) according to the manufacturer’s protocol.

### Estradiol and ACC treatment

For β-estradiol treatment, ten-day-old *HSFA7b-IOE* seedlings were incubated for 16 h with 10 μM β-estradiol or ethanol (0.1%, v/v; mock treatment). After incubation, seedlings were harvested for transcriptome analysis.

To check the sensitivity of Col-0 and *hsfa7a/b* mutants to ACC, seedlings were grown on 0.5 MS medium supplemented with or without different concentrations of ACC (0.25, 0.5, and 0.75 μM). Images were taken seven days after germination. To check the expression of *HSFA7a* and *HSFA7b* in response to ACC treatment, Col-0 seedlings were grown on 0.5 MS medium with or without 10 μM ACC for 10 days.

### Statistical analysis

Statistical significance was calculated using Student’s *t*-test: \**P* ≤ 0.05; \*\**P* ≤ 0.01; \*\*\**P* ≤0.001.

### Data availability

Sequencing data are available at the NCBI Sequencing Read Archive (SRA), BioProject ID PRJNA877651.

### AGI codes

*HSFA7a, AT3G51910; HSFA7b, AT3G63350; ERF1A, AT4G17500; EIN3, AT3G20770; TUB2, AT5G62690; SYD, AT2G28290*. Additional AGI codes are provided in the supplementary tables.

