## Supplemental Information for "Transcription factor HSFA7b controls ethylene signaling and meristem maintenance at the shoot apical meristem during thermomemory"

**This PDF file includes:**

Supplemental Figure 1 – 8

Supplemental Table 1 – 2

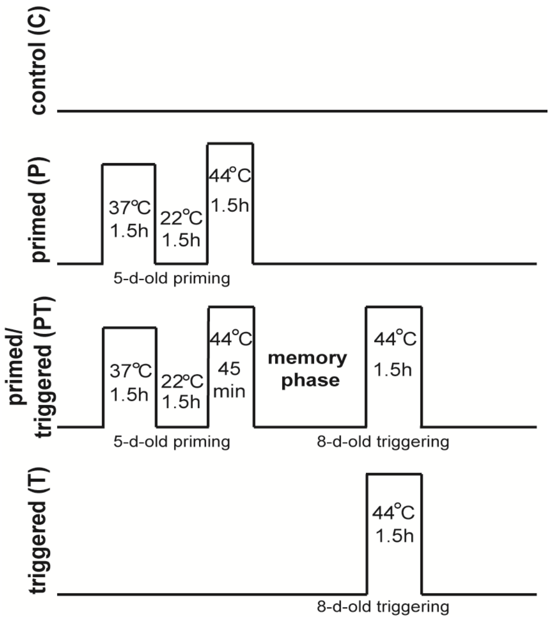


**Supplemental Figure 1. Thermomemory assay.**

```
HSFA7a             1 MMNPFLPEGCDPPPPPQPMEGLHENAPPPFLTKTFEMVDDPNTDHIVSWN     50
                      |:|.........|||.|||||.|..|.||||||||||.||||:||||||
HSFA7b             1 -MDPSSSSRARSMPPPVPMEGLQEAGPSPFLTKTFEMVGDPNTNHIVSWN     49

HSFA7a            51 RGGTSFVVWDLHSFSTILLPRHFKHSNFSSFIRQLNTYGFRKIEAERWEF    100
                     |||.||||||.||||..:||.:|||:|||||:||||||||||||||||||
HSFA7b            50 RGGISFVVWDPHSFSATILPLYFKHNNFSSFVRQLNTYGFRKIEAERWEF     99

HSFA7a           101 ANEEFLLGQRQLLKNIKRRNPFTPSSSP-----------SHD---ACNEL    136
                     .||.||:|||.|||:||||   |.||||           :||   ...:|
HSFA7b           100 MNEGFLMGQRDLLKSIKRR---TSSSSPPSLNYSQSQPEAHDPGVELPQL    146

HSFA7a           137 RREKQVLMMEIVSLRQQQQTTKSYIKAMEQRIEGTERKQRQMMSFLARAM    186
                     |.|:.||||||.:|||::|..:.|::||||||.|.|:|||.|||||.||:
HSFA7b           147 REERHVLMMEISTLRQEEQRARGYVQAMEQRINGAEKKQRHMMSFLRRAV    196

HSFA7a           187 QSPSFLHQLLKQ-RDKKIKELEDNESAKRKRGSSSMSELEVLALEMQGHG    235
                     ::||.|.|:.:| ||::...:.|....:.......:||||.|||||||:|
HSFA7b           197 ENPSLLQQIFEQKRDREEAAMIDQAGLIKMEEVEHLSELEALALEMQGYG    246

HSFA7a           236 KQRNMLEEEDHQLVVERELDDGFWEELL-----SDESLASTS--    272
                     :||.     |.   ||||||||||||||     |||..|:..
HSFA7b           247 RQRT-----DG---VERELDDGFWEELLMNNENSDEEEANVKQD    282
```

**Supplemental Figure 2. HSFA7a and HSFA7b protein sequence alignment.** The alignment was done using EMBOSS Needle (www.ebi.ac.uk/Tools/psa/emboss_needle).

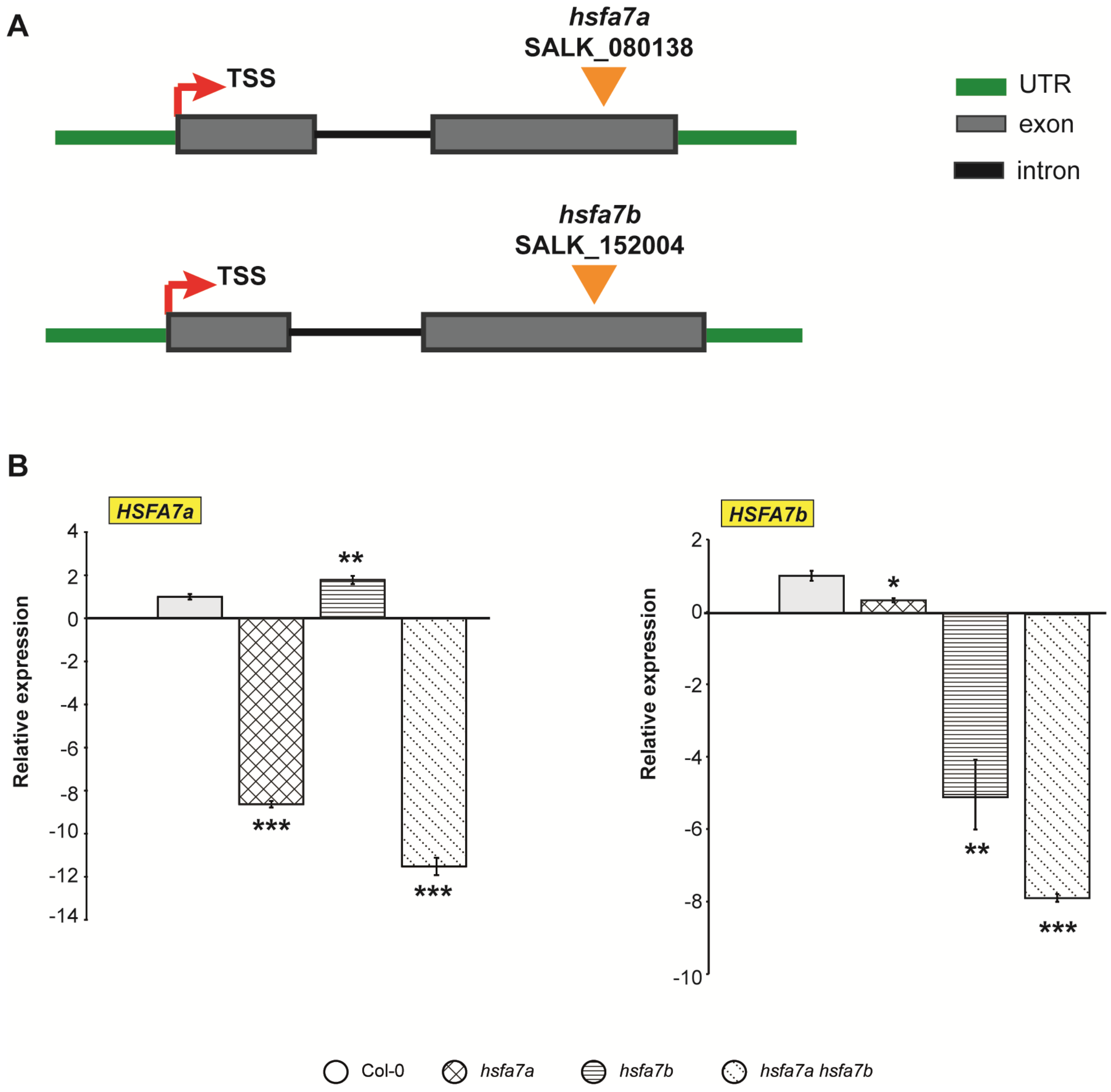


**Supplemental Figure 3. Characterization of *hsfa7a* and *hsfa7b* mutants.** **(A)** Schematic representation of the *hsfa7a* and *hsfa7b* mutants depicting the positions of T-DNA insertions. The green boxes represent the 5' and 3' UTRs, the black lines represent introns and the grey boxes represent exons. The red arrow represents the transcription start site (TSS). **(B)** Expression level of *HSFA7a* and *HSFA7b* measured in 5-day-old Col-0, *hsfa7a*, *hsfa7b* and *hsfa7a hsfa7b* mutant plants at 2 h after priming. Error bars indicate ± s.d. (n = 3). Asterisks indicate statistically significant differences (Student *t*-test: $*P \leq 0.05$; $**P \leq 0.01$; $***P \leq 0.001$) compared to Col-0 under same conditions.

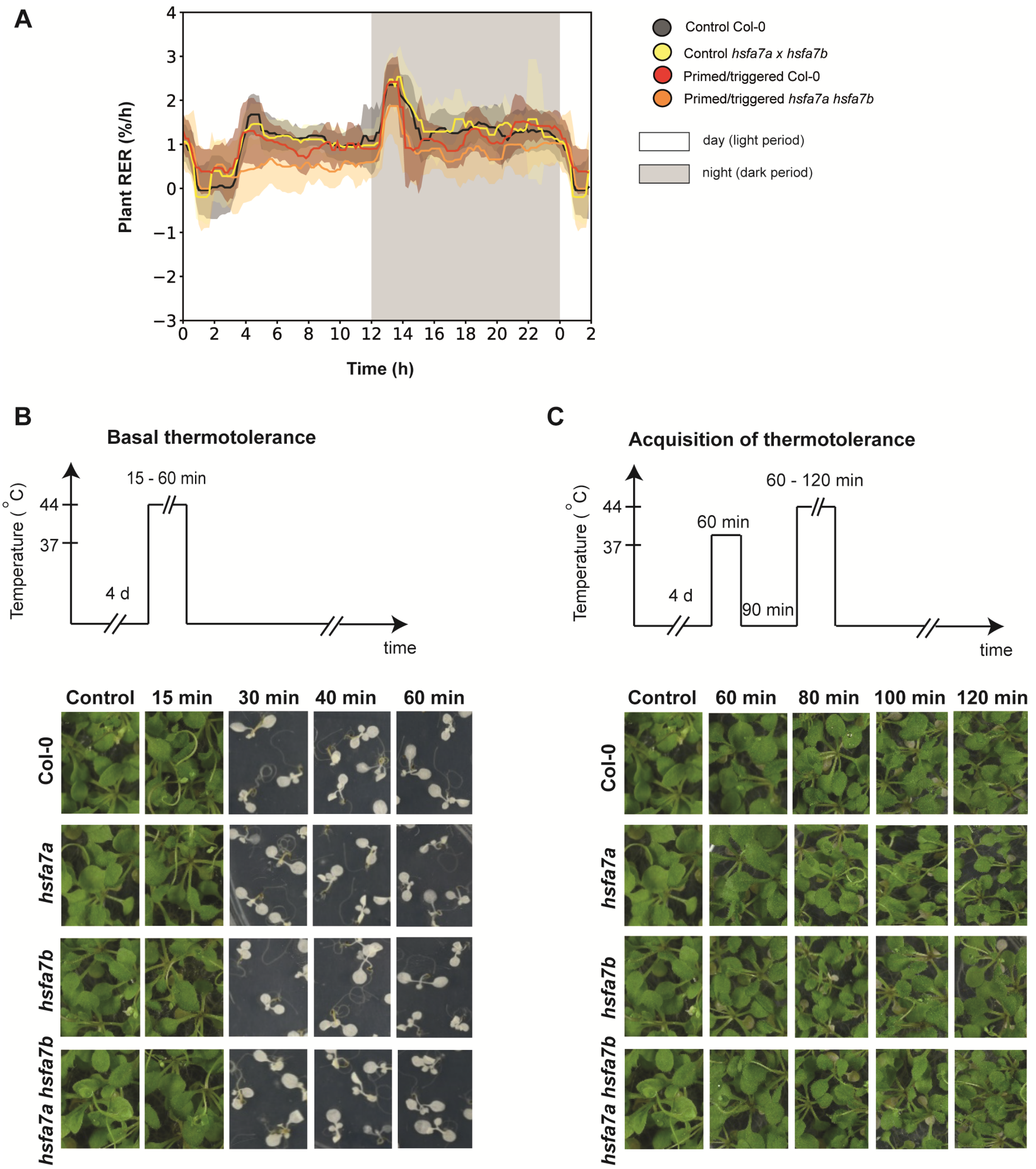


**Supplemental Figure 4. Analyses of diurnal relative expansion rate (RER), basal thermotolerance, and acquisition of thermotolerance. (A)** Time-resolved RER averaged over seven sequential 24-h periods of control (C) and primed/triggered (PT) Col-0 wild-type and *hsfa7a hsfa7b* mutant plants measured using an 3D imaging system. **(B)** Basal thermotolerance and **(C)** acquisition of thermotolerance of Col-0, *hsfa7a, hsfa7b,* and *hsfa7a hsfa7b* mutant plants.

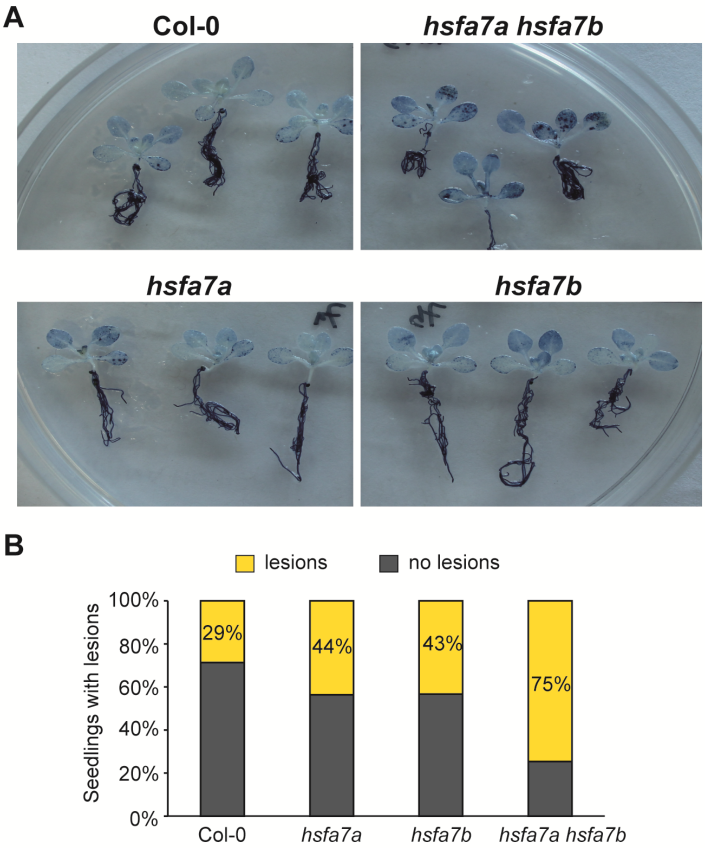


**Supplemental Figure 5. Thermopriming induces cell death in the *hsfa7a x hsfa7b* mutant. (A)** Trypan Blue staining for cell death in primed/triggered Col-0, *hsfa7a, hsfa7b,* and *hsfa7a hsfa7b* mutants at 24 h after triggering treatment. **(B)** Percentage of primed/triggered Col-0, *hsfa7a, hsfa7b,* and *hsfa7a hsfa7b* seedlings with lesions.

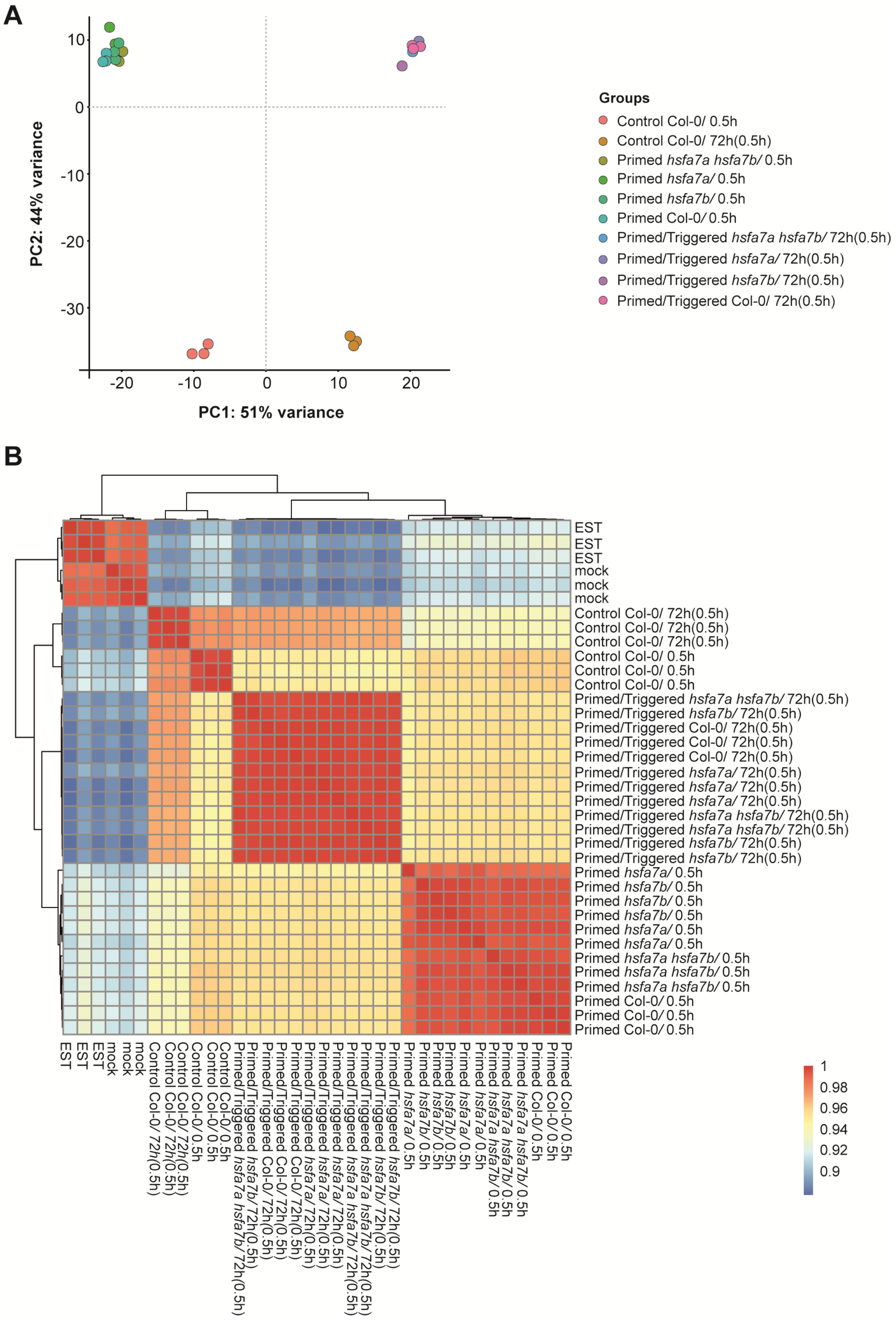


**Supplemental Fig. 6: Principal component analysis (PCA) and clustering heat map of gene expression analyzed in all samples. (A)** PCA describing the relationship between meristem samples of control Col-0, primed and primed/triggered Col-0, *hsfa7a, hsfa7b* single, and *hsfa7a hsfa7b* double mutants. **(B)** Heat map of the correlation matrix of all samples using pairwise Person correlation. Note that the clustering heat map revealed a weak outlier sample (Primed *hsfa7a*/ 0.5 h) which was removed for further analysis. Heat map and PCA plots were generated with normalized expression values generated by applying variance stabilizing transformation (VST) using DESeq2.

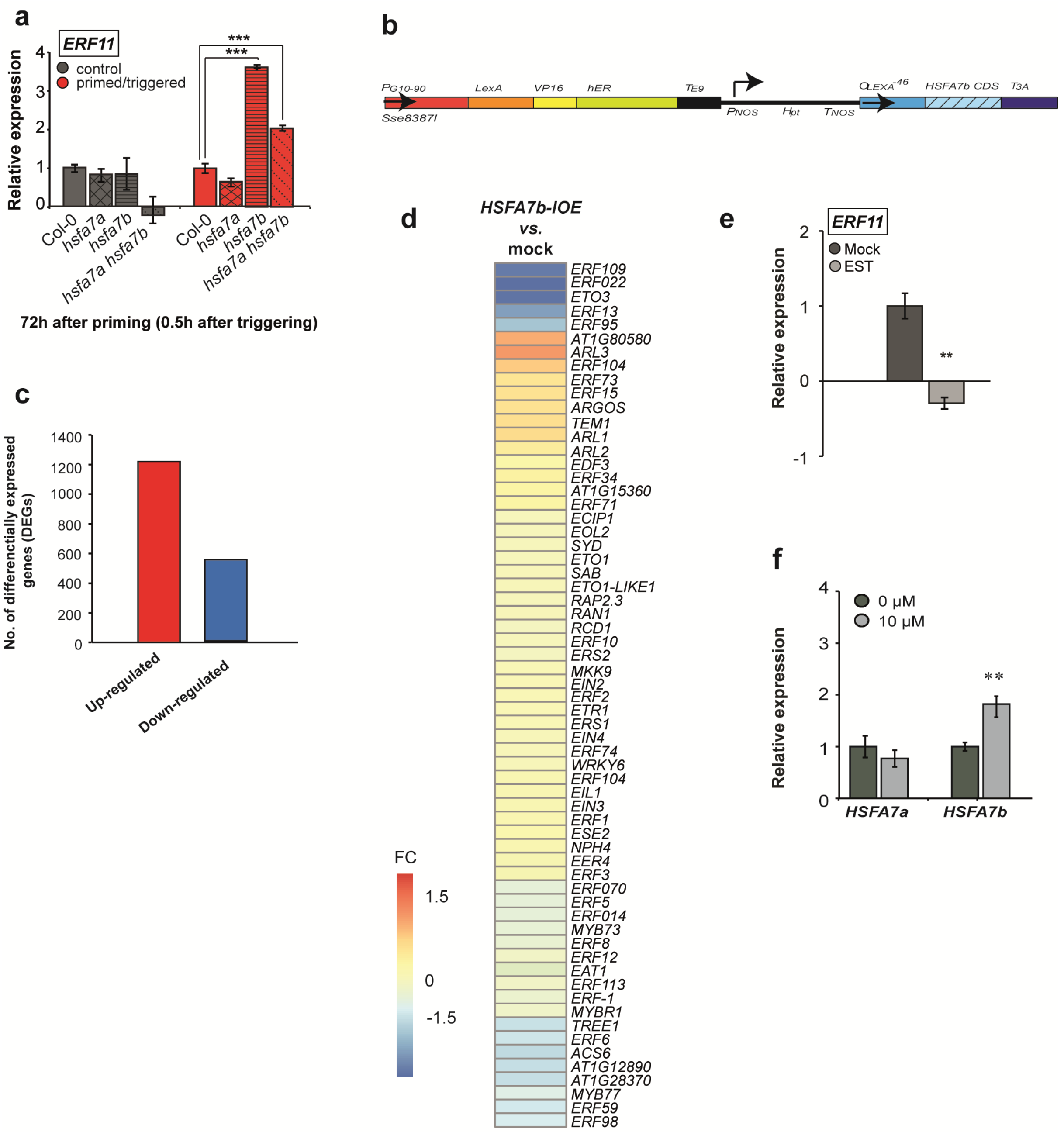


**Supplemental Fig. 7: HSFA7b controls ethylene response at the SAM during thermomemory. a,** Expression level of *ETHYLENE RESPONSE FACTOR 11* (*ERF11*) analyzed at the shoot apical meristem of control and primed/triggered Col-0, *hsfa7a, hsfa7b*, and *hsfa7a hsfa7b* plants at 0.5 h after triggering (72 h after priming). **b,** *HSFA7b-IOE* construct generated by inserting the *HSFA7b* coding sequence into the XVE vector (Zuo *et al.*, 2000). Black arrows indicate the direction of the transcription. Abbreviations: $P_{G10\text{-}90}$, a synthetic promoter controlling *XVE* expression; *LexA,* the transcriptional activation domain of *V16; hER,* human estrogen receptor; *TE9, rbcS E9* poly(A) addition sequence; *Pnos*, nopaline synthase promoter; *Hpt*, hygromycin phosphotransferase II coding sequence; *Tnos*, nopaline synthase poly(A) addition sequence; *OLexA*, eight copies of the *LexA* operator sequence; *HSFA7b CSD*, *HSFA7b* coding sequence; $T_{3A}$, *rbcs*S3A poly(A) addition sequence. **c,** Number of differentially expressed up- (red) and down-regulated genes in *HSFA7b-IOE*. Note that samples were harvested 16 h after β-estradiol induction**. d,** Heat map showing the log2 fold change (log2 FC) of the expression of ethylene-related up-regulated (red) or down-regulated (blue) genes in *HSFA7b-IOE* compared to mock-treated plants at 16 h after estradiol induction. **e,** Expression level of *HSFA7a* and *HSFA7b* in EST- and mock-treated *HSFA7b-IOE* plants. Error bars represent ± s.d. (*n* =

3). **f,** Expression level of *HSFA7a* and *HSFA7b* in response to ACC treatment as detected by qRT-PCR. Error bars represent ± s.d., ($n = 3$). Asterisks indicate statistically significant difference (Student´s *t*-test, $^{**}P < 0.01$, and $^{***}P < 0.001$) compared to Col-0 under the same condition (a,f) and compared to mock treatment (e).

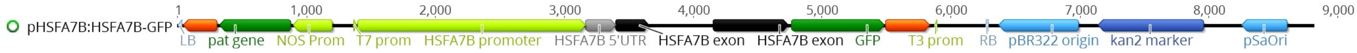


**Supplemental Fig. 8: *pHSF7b:HSFA7b:GFP* construct.** *pHSF7b:HSFA7b:GFP* was generated by cloning the *HSFA7b* promoter (2 kb) and coding sequence (without stop codon) fused to GFP using the pGREENII 0229 vector (Helens *et al.,* 2000).

**Supplemental Table 1. Number of significantly changed genes with $\log_2$ FC > |1.5|.**

| | 0.5 h after priming | 0.5 h after triggering (72 h after priming) |
|---|---|---|
| **DEGs: significant** | | |
| Col-0 (P) *vs*. Col-0 (C) | 12343 (6347↓; 5996↑) | NA |
| *hsfa7a* (P) *vs*. Col-0 (P) | 1406 (717↓; 689↑) | NA |
| *hsfa7b* (P) *vs*. Col-0 (P) | 3206 (1688↓;1518↑) | NA |
| *hsfa7a hsfa7b* (P) *vs*. Col-0 (P) | 810 (529↓; 281↑) | NA |
| Col-0 (PT) *vs*. Col-0 (C) | - | 9845 (5030↓; 4815↑) |
| *hsfa7a* (PT) *vs*. Col-0 (PT) | - | 274 (161↓;113↑) |
| *hsfa7b* (PT) *vs*. Col-0 (PT) | - | 153 (67↓; 86↑) |
| *hsfa7a hsfa7b* (PT) *vs*. Col-0 (PT) | - | 1062 (518↓; 544↑) |
| **DEGs: significant and $\log_2$FC > \|1.5\|** | | |
| Col-0 (P) *vs*. Col-0 (C) | 8001 (3758↓; 4243↑) | NA |
| *hsfa7a* (P) *vs*. Col-0 (P) | 668 (284↓; 384↑) | NA |
| *hsfa7b* (P) *vs*. Col-0 (P) | 1042 (481↓; 561↑) | NA |
| *hsfa7a hsfa7b* (P) *vs*. Col-0 (P) | 204 (162↓; 42↑) | NA |
| Col-0 (PT) *vs*. Col-0 (C) | - | 4366 (1576 ↓; 2790↑) |
| *hsfa7a* (PT) *vs*. Col-0 (PT) | - | 63 (26↓; 37↑) |
| *hsfa7b* (PT) *vs*. Col-0 (PT) | - | 53 (21↓; 32↑) |
| *hsfa7a hsfa7b* (PT) *vs*. Col-0 (PT) | - | 252 (26↓; 226↑) |

Abbreviations: C, control; P, primed; PT, primed and triggered; T, triggered; NA, not applicable. Downward directed arrows (↓) indicate downregulated genes; upward directed arrows (↑) indicate upregulated genes.

**Supplemental Table 2. Oligonucleotides used in this study.**

| Gene (AGI) | Oligonucleotide | Sequence (5'→3') |
|---|---|---|
| **Oligonucleotides used for cloning** | | |
| ***ERF1A*** | ERF1A_F | ATGTCGATGACGGCGGATTC |
| *AT4G17500* | ERF1A_R | TTATAAAACCAATAAACGATCGCC |
| ***HSFA7a*** | HSFA7a_F | ATGATGAACCCGTTTCTCCC |
| *AT3G51910* | HSFA7a_R | TTAGGAGGTGGAAGCCAAACTC |
| ***HSFA7b*** | HSFA7b_F | ATGGACCCGTCGTCAAGCTCC |
| *AT3G63350* | HSFA7b_R | CTAATCTTGCTTCACATTCGC |
| ***HSP17.8*** | HSP17.8_F | ATGTCGCTTATTCCAAGCTTC |
| *AT1G07400* | HSP17.8_R | TTAGCCAGAGATATCAATAGAC |
| ***WUS*** | WUS_F | ATGGAGCCGCCACAGCATCAGC |
| *AT2G17950* | WUS_R | CTAGTTCAGACGTAGCTCAAGAG |
| ***STM*** | STM_F | ATGGAGAGTGGTTCCAACAGC |
| *AT1G62360* | STM_R | TCAAAGCATGGTGGAGGAGATG |
| **Oligonucleotides used for qRT-PCR** | | |
| ***HSFA7a*** | HSFA7a_qRT_F | ACCACCACCACAACCAATGGAG |
| *AT3G51910* | HSFA7a_qRT_R | TCTTGGTCAGAAATGGAGGTGGAG |
| ***HSFA7b*** | HSFA7b_qRT_F | ATGGAGGGATTGCAGGAAGCAG |
| *AT3G63350* | HSFA7b_qRT_R | TGGATCACCAACCATCTCGAACG |
| ***HSFA2a*** | HSFA2a_qRT_F | GCAGCGTTGGATGTGAAAGTGG |
| *AT2G26150* | HSFA2a_qRT_R | TTGGCTGTCCCAATCCAAAGGC |
| ***WUS*** | WUS_qRT_F | AGAAGAAGAATGTGGTGGCGATGC |
| *AT2G17950* | WUS_qRT_R | AGACGTAGCTCAAGAGAAGCGCAA |
| ***STM*** | STM_qRT_F | TCGACTTCTTCCTCGGATGACCCA |
| *AT1G62360* | STM_qRT_R | TCTCCGGTTATGGAGAGACAGCAA |
| ***ERF1A*** | ERF1A_qRT_F | TTGCGGCGGAGATTAGAGAC |
| *AT4G17500* | ERF1A_qRT_R | ATTCAACAAAGCGCGGGAAC |
| ***ERF11*** | ERF11_qRT_F | AGGCTGGGATGATGGTGTTC |
| *AT1G28370* | ERF11_qRT_R | GTTCTCAGGTGGAGGAGGGA |
| ***ERF104*** | ERF104_qRT_F | AGAGAGGCACTACAGGGGAG |
| *AT5G61600* | ERF104_qRT_R | GTGTCGTAAGTCCCAAGCCA |
| ***EIN2*** | EIN2_qRT_F | AATGACACCGTGCTTTTGCC |
| *AT5G03280* | EIN2_qRT_R | TGACTGCGGTTGTGCATTTG |
| ***EIN3*** | EIN3_qRT_F | TGTCTGGTGGAAGTTGCTCG |
| *AT3G20770* | EIN3_qRT_R | ATTCCGAGTTTCCTGCTGGG |
| ***EIL1*** | EIL1_qRT_F | AAGCAACCAAACGCCTCCTA |
| *AT2G27050* | EIL1_qRT_R | TTAACCCCGTTGTTCGTCCC |
| ***SYD*** | SYD_qRT_F | CAGTGGGAGGTATGGACGTT |
| *AT2G28290* | SYD_qRT_R | CACCGTCAAGCCTGTCTGAT |
| ***TUB2*** | TUB_qRT_F | GAGCCTTACAACGCTACTCTGTCTGTC |

| | | |
|---|---|---|
| *AT5G62690* | TUB_qRT_R | ACACCAGACATAGTAGCAGAAATCAAG |
| | **Oligonucleotides used for ChIP-qPCR** | |
| ***EIN3*** *AT3G20770* | EIN3_F1<br>EIN3_R1 | TCCATTCAAAGGGACAGGGA<br>AGACTGATGGAAATAAAGGCGGA |
| | EIN3_F2<br>EIN3_R2 | CTAGCTGAGCATGTAGAACAGGT<br>GTAGTCCACCTGAAACCACCA |
| | EIN3_NF<br>EIN3_NR | CACGCAATGACGCAAATCCT<br>CACTCGACCTCGTGAACACA |
| ***SYD*** *AT2G28290* | SYD_F1<br>SYD_R1 | ATCAAGAACAACCGCGACCT<br>TCGCTGGTTTTTGGTTTCCT |
| | SYD _F2<br>SYD_R2 | ACTTCACGCACTTGTGGTAGAT<br>AATCTTTCACTGCCAAGGGT |
| | SYD _NF<br>SYD _NR | AGGATTTCCTGAGTTTTTACATAGC<br>GCTGCCTCCATTCATAGCCT |
| | **Oligonucleotides used for genotyping** | |
| *hsfa7a* (SALK_080138) | hsfa7a_LP<br>hsfa7a_RP<br>LBb1.3 | GTTCCAGAAGCAAGTTTCGTG<br>TTGCTCACTCATGTGGACTTG<br>ATTTTGCCGATTTCGGAAC |
| *hsfa7b* (SALK_152004) | hsfa7b_LP<br>hsfa7b_RP<br>LBb1.3 | TTCTTCGCAAGTTCTGGAAAC<br>TCCCATTTTATAAGATTTTCAAGC<br>ATTTTGCCGATTTCGGAAC |
| *erf1a* (SALK_036267) | erf1a_LP<br>erf1a_RP<br>LBb1.3 | CGTTCCTAACCAAACCCTAGC<br>TCCTACTCTTCTCCCTGCTCC<br>ATTTTGCCGATTTCGGAAC |
| *erf11* (SALK_116053) | erf11_LP<br>erf11_RP<br>LBb1.3 | CCACACGTCGTCCTTCATATC<br>TGCAAAGCCTAAAATTAAAAACG<br>ATTTTGCCGATTTCGGAAC |
